# Pharmacokinetics of long-term low-dose oral rapamycin in four healthy middle-aged companion dogs

**DOI:** 10.1101/2021.01.20.427425

**Authors:** Jeremy B. Evans, Ashley J Morrison, Martin A. Javors, Marisa Lopez-Cruzan, Daniel E.L. Promislow, Matt Kaeberlein, Kate E. Creevy

## Abstract

**Objective:** To determine the blood concentration and pharmacokinetic parameters of rapamycin in companion dogs following long-term, low-dose oral administration of rapamycin.

**Animals:** Four healthy, middle-aged, medium-to-large breed privately owned dogs participated.

**Procedures:** All dogs had been receiving oral rapamycin at a dose of 0.025 mg/kg on Monday, Wednesday, and Friday mornings for at least one month. An initial blood sample was collected prior to morning rapamycin administration, and samples were collected at 1, 2, 6, and 24 hours after rapamycin was given. Blood samples were transferred to blood spot collection cards, air-dried and stored at −80°C. Rapamycin concentrations were determined via HPLC/MS. All blood collections occurred on Wednesdays, so that the previous dose of rapamycin had taken place 48 hours prior to blood collection.

**Results:** For all dogs, rapamycin T_max_ was 2 hours. Median C_max_ was 1.47 ng/ml (0.912 – 2.13), and the median AUC_0-last_ was 15.7 ng*hr/mL (1.30 – 36.3). Due to sample size and timing, the only estimates related to elimination rate reported are for mean residence time with a median of 4.70 hrs (0.90 – 7.30).

**Conclusions and Clinical Relevance:** A 0.025 mg/kg oral dose of rapamycin, administered three times a week, resulted in concentrations of rapamycin in the blood capable of being measured in ng/ml.

## Introduction

Rapamycin (sirolimus), an FDA-approved natural macrolide that is produced by *Streptomyces hygroscopicus,* is an inhibitor of the mechanistic target of rapamycin (mTOR). As part of the phosphatidylinositol kinase-related kinase (PIKK) family, mTOR is a serine/threonine kinase that is present within the mTOR complex 1 (mTORC1) and mTOR complex 2 (mTORC2). [1–3] The mTORC1 pathway is crucial for cell survival and drives cell growth and protein synthesis based in part on nutrient availability. In this regard, the activity of the nutrient/stress-sensing mTORC1 is increased during periods of nutrient abundance, and is conversely inhibited when under nutrient stress, such as reduced intracellular ATP or reduced amino acid availability. [1, 4–7] A similar inhibition of mTORC1 occurs under conditions of dietary restriction or intermittent fasting. [8] With rapamycin, inhibition of mTOR is due to the binding of rapamycin to the FK506 binding protein (FKBP12), which allosterically inhibits mTORC1. [9–11] This results in several downstream consequences including reduced translation of proteins that play a key role in the G1 to S transition of the cell cycle and activation of the autophagic degradation pathway. [1, 12, 13]

In humans, oral rapamycin is often used in immunosuppressive protocols in transplant recipients to prevent organ rejection.[14–16] In some cases, due to the mTOR pathway’s close association with insulin signaling, rapamycin treatment has resulted in derangements in glucose homeostasis.[17, 18] However, multiple studies have documented potential benefits of rapamycin treatment in the context of normative aging. For instance, rapamycin-mediated inhibition of mTORC1 has been shown to reduce age-related cancers [19, 20] and age-related declines in cognitive function [21, 22], and to improve heart function [23–25], immune function [26], kidney function [27], oral health [28, 29], intestinal function and gut dysbiosis [30, 31], and ovarian function [32] in aging mice. In both dogs and mice, rapamycin therapy improves cardiac function and can potentially treat hepatic glycogen storage disease.[4, 24, 25, 33–35] Inhibition of mTOR also results in decreased smooth-muscle cell migration and proliferation within coronary arteries, with rapamycin-coated coronary stents significantly reducing arterial stenosis following stent placement.[36] Still, one of the most striking findings is that rapamycin administration has been associated with a significant increase in subject lifespan, a result observed in multiple model organisms, including yeast [37], fruit flies [38], nematodes [39], and mice.[40–46]

In humans, oral rapamycin is absorbed rapidly with peak concentrations occurring within one to three hours depending on dosing protocol.[47, 48] In the bloodstream, the vast majority of rapamycin is distributed within red blood cells, and co-administration with a high fat meal can increase the oral bioavailability and AUC by up to 35% while decreasing the maximum blood concentration.[49, 50] In dogs, Larson *et al.* investigated rapamycin pharmacokinetics following a single oral dose of 0.1 mg/kg or a five-day course of this treatment.[51] In the single-dose protocol, rapamycin concentration peaked twice, at two- and six-hours post-administration, but this pattern was not noted when the dogs were treated on five consecutive days. These low-dose rapamycin treatments resulted in peak blood concentrations similar to those that displayed anti-tumorigenic properties in mice, with 8.39 ng/ml and 5.49 ng/ml measured in the single and consecutive dose protocols, respectively. In both instances, the time to reach maximum blood concentration was approximately three to four hours.[51] A different study found that intramuscular injections of rapamycin doses ranging from 0.01-0.08 mg/kg, either given once or daily over a seven-day treatment schedule, also resulted in measurable and clinically relevant blood concentrations of rapamycin. However, the time to maximum blood concentration in this study ranged from 2 to 48 hours, implying significantly greater variability with this route.[52] The goal of the study reported here was to determine the pharmacokinetics of oral low-dose rapamycin after long-term administration (two to six months) to healthy middle-aged dogs.

## Materials and Methods

### Study Participants

Seven healthy, middle-aged to senior dogs (three spayed females, four castrated males) between the ages of six to ten years old and weighing between 18.2-36.4 kg (40-80 lbs) were enrolled. All participating dogs had previously been recruited based on specified criteria into a 12-month prospective, double-blinded, placebo-controlled randomized clinical trial in which dogs received rapamycin or placebo three times per week for six months. Therefore, dogs in this ancillary study had been receiving treatment (either placebo or rapamycin) for durations ranging from one to six months based upon their date of enrollment into the clinical trial. At the time of this study, investigators remained masked to treatment assignment and collected blood samples from a total of seven dogs. Three of these dogs were later determined to have been in the placebo group, such that results from four rapamycin-treated dogs are reported here.

### Drug and placebo

Both 0.5 mg tablets (Cadlia Health Care, Zydus, Ahmedabad, India) and 1.0 mg tablets (Dr. Reddy’s Laboratories, Princeton, New Jersey; Greenstone LLC, Pfizer Inc.) of rapamycin were used. Rapamycin was administered to dogs at a dose of approximately 0.025 mg/kg, using 0.25 mg increments. As such, the dose of rapamycin was 0.50 mg for dogs between 18.2 and 23.0 kg (40-50 lbs), 0.75 mg for dogs between 23.1 and 30.0 kg (51-66 lbs), and 1.0 mg for dogs between 30.1 and 36.4 kg (67-80 lbs). To mask the treatment, a combination of tablets comprising the total daily dose was placed into gelatin capsules for dogs in the treatment group. Placebo capsules (Professional Compounding Centers of America; PCCA) contained lactose.

### Experimental design

All dogs were fasted overnight and underwent an initial blood draw at hour 0. Approximately four ml of blood were obtained via jugular venipuncture at every time point. After the initial blood draw at 0 hour, dogs were offered their normal breakfast and given either placebo or rapamycin. Additional blood samples were obtained via the same method from each dog at 1, 2, 6, and 24 hours after treatment administration. All blood samples were promptly transferred to K-EDTA blood storage tubes. Within an hour of each time point, up to 50 μL of whole blood were pipetted from the storage tube onto two spots of a blood spot card (Whatman 903 DBS (dried blood spot) collection cards; MilliporeSigma, St. Louis, MO). Two spot cards with two spots each were prepared for each dog. The spot cards were allowed to dry at room temperature for one hour and stored at −80°C in a sealed plastic bag with a desiccant packet (0.5 gram silica gel packets, Intertek Packaging, Orchard Park NY 14127) before being shipped for analysis.

All procedures for this study were reviewed and approved by the TAMU Institutional Animal Care and Use Committee (IACUC 2018-0299 CA). Because these were client-owned animals, an IRB determination was requested, and this study was found not to be Human Subjects Research (HSR).

### Analysis of Rapamycin Concentration

#### Measurement of Rapamycin Using HPLC/MS/MS

Rapamycin concentrations were determined via HPLC/MS/MS (Biological Psychiatry Analytical Lab [BPAL], University of Texas Health Science, San Antonio, TX). Rapamycin, ascomycin and all reagents were purchased from Sigma Chemical Company (St. Louis, MO). Milli-Q water was used for preparation of all solutions. The HPLC system consisted of a Shimadzu SIL 20A HT autosampler, LC-20AD pumps (2), and an AB Sciex API 3200 tandem mass spectrometer with turbo ion spray. The LC analytical column was a Grace Alltima C18 (4.6 x 150 mm, 5 μm) purchased from Alltech (Deerfield, IL) and was maintained at 25°C during the chromatographic runs using a Shimadzu CT-20A column oven. Mobile phase A contained 10 mM ammonium formate and 0.1% formic acid dissolved in 100% HPLC grade methanol. Mobile phase B contained 10 mM ammonium formate and 0.1% formic acid dissolved in 90% HPLC grade methanol. The flow rate of the mobile phase was 0.5 ml/min. Rapamycin was eluted with a gradient. Mobile phase gradient: 0-0.1 min, 100% B; 0.1-4 min, linear gradient to 0% B; 4.0-5.0, 0% B; 5-10 min, 100% B. The rapamycin transition was 931.6 m/z to 864.5 m/z, which was used for quantification. The internal standard (ascomycin) transition was 809.6 m/z to 756.6 m/z.

Rapamycin and ascomycin super stock solutions were prepared in methanol at a concentration of 1 mg/ml and stored in aliquots at −80°C. A working stock solution was prepared each day from the super stock solution at a concentration of 10 μg/ml and used to spike the calibrators. Calibration samples were pipetted on spot cards, dried, and then used to quantify rapamycin in the unknown samples.

Spiked calibration samples were prepared at concentrations of 0, 1.56, 6.25, 25.0 and 100 ng/ml. Two blood spots, each containing 50 μl, were cut into four segments and placed in 10 × 50 mm polypropylene tubes with push caps. One ml of mobile phase A and 10 μl of a 0.5 μg/ml solution of ascomycin were added to each tube. The tubes were capped, vortexed vigorously for 30 sec and placed on a shaker Model E6010 from Eberbach Corporation (Ann Arbor, Michigan) for 30 min. After shaking, the tubes were centrifuged for 10 min at 5,000 g. The supernatants were carefully poured or pipetted into 10 × 75 mm glass tubes and dried to residue under a nitrogen stream. The residues were then redissolved in 100 μl of mobile phase A and transferred to autosampler vials. Then, 50 μl were injected into the HPLC/MS/MS system. The ratios of rapamycin peak areas to ascomycin peak areas of unknown samples were compared against the linear regression of the ratios of the calibration samples to quantify rapamycin. The concentration of rapamycin was expressed as ng/ml of blood.

#### Pharmacokinetic Analysis

Noncompartmental analysis using industry standard software (Certera Phoenix WinNonLin 8.2.0.4383, Princeton, NJ) was attempted using the blood concentrations from individual dogs to estimate pharmacokinetic parameters. Attempts were made to estimate or calculate the following parameters: T_max_: time of maximum observed blood concentration; C_max_: maximum observed blood concentration; Az: terminal elimination rate; t_1/2γz_: apparent terminal half-life; AUC_0.last_: area of the curve from time zero to last measurable time; AUC_0-inf_: area under the curve from time zero extrapolated to infinite time; AUC % Extrap: area under the curve extrapolated from time zero to infinity as a percent of total AUC; AUMC_0-obs_ = area under the moment curve from time zero to last observed concentration; AUMC_0-inf_ = area under the moment curve from time zero extrapolated to infinity; MRT_0-obs_ = mean resident time estimated using time zero to last observed concentrations, calculated as AUMC_0-obs_ /AUC_0-obs_; MRT_0-inf_ = mean residence time estimated using time zero to infinity, calculated as AUMC_0-inf_ /AUC_0-inf_.

## Results

### Study Participants

The participant demographics are displayed in Table 1. A total of seven dogs were sampled. Of these, four dogs (three spayed females, one castrated male) were found to be receiving rapamycin after the clinical trial was unblinded. There were three purebred (Labrador Retriever, Australian Shepherd, Pit Bull Terrier) and one mixed breed dog represented. The median age of this group of dogs was 9.0 years (range: 8.8 – 11 years) and they were all medium to large breeds (median weight: 27.0 kg, range: 20.2 – 34.0 kg). As dogs had been previously enrolled at different dates into the clinical trial, the treatment period prior this this study varied. The range of treatment period prior to this study was one to five months; two of the four dogs had been receiving treatment for one month. Each dog had been prescribed a dose of rapamycin according to its weight, with one dog receiving 0.50 mg, two dogs receiving 0.75 mg, and one dog receiving 1.0 mg. While all dogs were offered breakfast at the time of medication administration, only one dog (#4) ate.

**Table 1.** Participant demographics.

| **Dog**<br>**#** | **Age**<br>**(years)** | **Breed** | **Sex** | **Weight**<br>**(kg)** | **Treatment** | **Treatment**<br>**Duration**<br>**(months)** |
| --- | --- | --- | --- | --- | --- | --- |
| **1** | 7.8 | Border Collie | FS | 19.0 | Placebo | 5.0 |
| **2** | 9.2 | Labrador<br>Retriever | MC | 34.0 | 1.0 mg<br>rapamycin | 1.0 |
| **3** | 8.8 | Australian<br>Shepherd | FS | 26.0 | 0.75 mg<br>rapamycin | 1.0 |
| **4<sup>a</sup>** | 9.8 | Pit bull terrier | FS | 28.0 | 0.75 mg<br>rapamycin | 5.0 |
| **5** | 10.8 | Australian<br>Shepherd | MC | 26.2 | Placebo | 4.0 |
| **6** | 11.0 | Mixed breed | FS | 20.2 | 0.50 mg<br>rapamycin | 3.0 |
| **7** | 8.5 | Border Collie | MC | 29.0 | Placebo | 5.5 |

<sup>a</sup>Only participant who received rapamycin with a meal

### Blood Rapamycin Concentrations

All four dogs had received rapamycin approximately 48 hours prior to baseline collection for this study. No dogs had detectable levels of rapamycin in their blood at baseline (time 0), just prior to their next scheduled administration of rapamycin. No placebo dogs had detectable levels of rapamycin in their blood at any time. Following administration of rapamycin, all four treated dogs (#2, 3, 4, and 6) had measurable rapamycin blood concentrations that were detectable by HPLC/MS/MS (Table 2).

**Table 2.** Blood concentration of rapamycin (ng/ml) in four dogs following a 0.025 mg/kg oral dose of rapamycin.

| Time (hrs) | Dog #2 | Dog #3 | Dog #4 | Dog #6 | Median |
| --- | --- | --- | --- | --- | --- |
| 0 | 0 | 0 | 0 | 0 | 0 |
| 1 | 1.07 | 1.15 | 0.884 | 0.861 | 0.988 |
| 2 | 1.08 | 1.86 | 2.13 | 0.912 | 2.03 |
| 6 | 0.933 | 1.10 | 1.80 | 0 | 1.02 |
| 24 | 0 | 0.875 | 1.14 | 0 | 0.438 |

By one hour post-administration, rapamycin concentrations increased for all four dogs (Figure 1), with the median rapamycin concentration one_hour post-administration at 0.980 ng/ml (0.861 – 1.15). At two hours post-administration, concentrations ranged from 0.912 to 2.13 ng/ml (median = 1.47). Following this two-hour time point, blood concentrations progressively decreased in all dogs, with the median rapamycin concentrations measured at six and 24 hours being 1.02 ng/ml (0.000 – 1.80) and 0.440 ng/ml (0.000 – 1.14). By six hours postadministration, dog #6 no longer had detectable rapamycin in circulation, and by 24 hours postadministration, dog #2 also no longer had detectable rapamycin in his blood.

**Figure 1:**
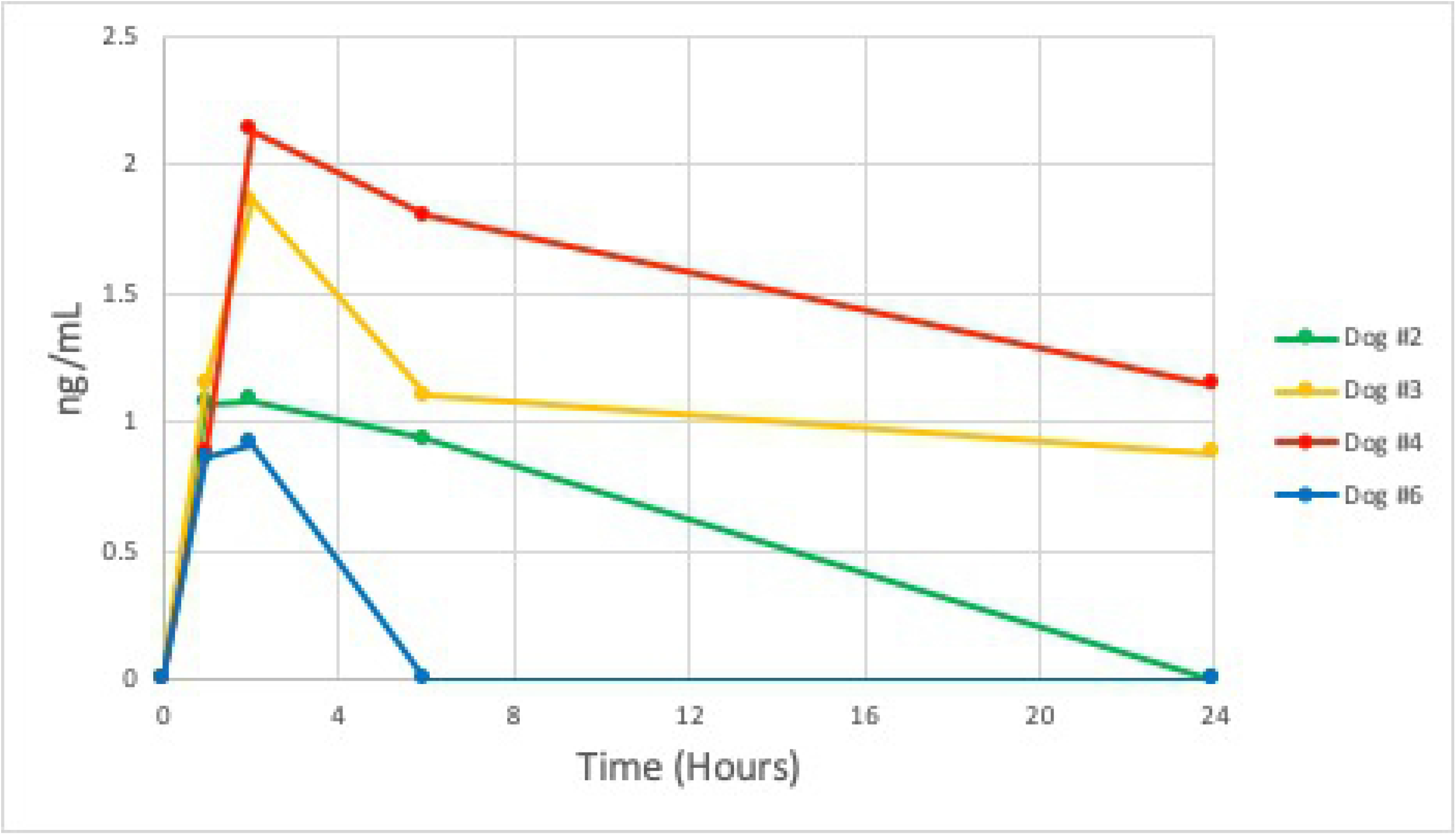
Blood concentration (ng/ml) of rapamycin in four dogs receiving long-term low-dose rapamycin therapy before (0), and 1, 2, 6, and 24 hours after oral administration.

### Pharmacokinetic Parameters

Estimates of pharmacokinetic parameters are presented in Table 3. For all dogs, T_max_ was 2 hours, and the median C_max_ was 1.47 ng/ml (0.91 – 2.1). Due to the sparse sampling and lack of sufficient samples for accurate nonlinear regression of the terminal phase drug concentrations, apparent elimination half-life, AUC extrapolated to infinity, and AUMC extrapolated to infinity could not be calculated. Therefore, the only estimates related to elimination rate reported are for MRT. The median MRT was 6.8 hrs (1.3 – 10). The median AUC_0-last_ was 15.7 ng*hr/mL (1.30 – 36.3).

**Table 3:** Pharmacokinetic parameters of rapamycin in four dogs following a 0.025 mg/kg oral dose.

| **Dog#** | **Units** | **2** | **3** | **4** | **6** | **Median** |
| --- | --- | --- | --- | --- | --- | --- |
| **$T_{\max}$** | **hr** | 2 | 2 | 2 | 2 | 2 |
| **$C_{\max}$** | **ng/ml** | 1.1 | 1.9 | 2.1 | 0.9 | 1.47 |
| **$AUC_{0-\text{last}}$** | **hr\*ng/ml** | 5.6 | 25.8 | 36.3 | 1.3 | 15.7 |
| **$AUMC_{0-\text{last}}$** | **hr\*ng\* ng/ml** | 17.7 | 272.1 | 376.6 | 1.8 | 144.9 |
| **$MRT_{\text{last}}$** | **hr** | 3.1 | 10.6 | 10.4 | 1.3 | 6.8 |

$T_{\max}$ : time of maximum observed blood concentration; $C_{\max}$ : maximum observed blood concentration; $AUC_{0-\text{last}}$ : area of the curve from time zero to last measurable time; $AUMC_{0-\text{last}}$ : area under the moment curve from time zero to last measurable time; $MRT_{\text{last}}$ : mean residence time calculated from $AUMC_{0-\text{last}} / AUC_{0-\text{last}}$ .

## Discussion

The results of this study indicate that low-dose (0.025 mg/kg, three times per week for one to five months) oral rapamycin, administered to healthy, middle-aged dogs, resulted in low but measurable concentrations of rapamycin in the blood. Similar concentrations have resulted in reduced mTOR activity in canine tumor cells.[52] Though rapamycin has previously been administered to dogs in other studies, this study is unique in the duration of treatment with a lower dose than previously described. Prior to this report, the longest published study of rapamycin in companion dogs was 10 weeks (2.5 months), during which Urfer *et al.* employed a dosing schedule of three days a week, on Monday, Wednesday, and Friday mornings. However, the doses used in that study were higher (either 0.05 mg/kg or 0.1 mg/kg). Additionally, no analysis of drug concentrations was performed. [33] Larson, *et al* [51] described pharmacokinetic analysis of oral rapamycin in companion dogs treated with 0.1 mg/kg administered either once or on five consecutive days. By comparison, the lower-dose, three-day-a-week regimen utilized in this study is markedly different. This three-times-weekly dosing schedule is intended to be employed in a nationwide study, with a much larger cohort of dogs who will receive rapamycin for three years. Because of this, investigation into the pharmacokinetics of rapamycin administration using this novel dosing schedule was warranted.

Not surprisingly, some of the pharmacokinetic parameters obtained from this study differed from those obtained by Larson, *et al*.[51] In the current study, the median C_max_ was 1.47 ng/ml, with the highest rapamycin blood concentration measured being 2.13 ng/ml. By contrast, maximum blood concentrations following a 0.1 mg/kg oral dose of rapamycin were greater, with a single oral dose or five consecutive daily doses resulting in mean concentrations of 8.39 ng/ml and 5.49 ng/ml, respectively.[51] On the other hand, despite administering rapamycin via intramuscular injections rather than orally, Paoloni *et al.* reported a similar C_max_ to this study (median: 1.69 ng/ml, range: 1.21-1.82 ng/ml) in dogs receiving a 0.02 mg/kg dose on eight consecutive days.[52] This might indicate that in dogs, maximum blood concentrations of rapamycin are more affected by the dose utilized, rather than the frequency or route of administration. The median Cmax in this study is significantly lower than that considered therapeutic for preventing organ transplant rejection in humans, where a blood concentration of 8-15 ng/ml is often desired.[14, 53] This is intentional, as our goal was to use a dose that results in measurable blood concentrations, but not with the objective of causing immunosuppression. The T_max_ reported in our study (two hours) is more similar to that reported by Larson *et al.* (3.3 ± 2.5 hours after one dose, and 4.5 ± 1.0 hours after five consecutive doses) compared to that observed after intramuscular (IM) injection (up to 48 hours), indicating faster absorption with oral administration.[51, 52]

The median MRT was 6.8 hours. Because AUC could not be extrapolated to infinity using regression analysis, it is difficult to assess the sufficiency of sampling times. Typically, one would look at area under the curve extrapolated from time zero to infinity as a percent of total AUC to evaluate whether samples were collected for long enough after drug dosing, but there were not enough non-zero sampling times after C_max_ to regress appropriately. Due to this, T_1/2_ was unable to be directly calculated, and MRT was reported.

In a previous study utilizing rapamycin, a mean half-life of 38.7 hours was found by Larson *et al.* following a single dose of 0.1 mg/kg orally. This was far lower than the T_1/2_ of 99 hours observed following five consecutive 0.1 mg/kg oral doses in that same study.[51]. It should be noted that, similar to the variable T_1/2_ calculated amongst drug recipients in both previous canine rapamycin pharmacokinetic studies (oral and IM administration), our MRT was not consistent between recipients. While two of the four dogs had an estimated MRT within 0.2 hours (10.4 & 10.6 hrs), the MRT of the other two dogs was significantly less (3.1 & 1.3 hrs).[51, 52] Whether the MRT calculated in this study is due to the low rapamycin dose utilized, the intermittent dosing schedule, the duration of administration, or a combination of these factors, is unknown. Additionally, because none of our dogs had measurable levels of rapamycin in their blood after 48 hours (that is, at time zero in our study), this may indicate that our dosing regimen was more akin to intermittent pulse-therapy as opposed to a long-term, continuous treatment. At this time, the benefit or significance of this is unknown.

This study is not without limitations. All four dogs utilized were already participating in a 12-month prospective study investigating rapamycin administration, and therefore, the duration of treatment prior to pharmacokinetic analysis was variable. Additionally, the sample size is small, attributable to the strict standards dogs must have met to be enrolled in the previously mentioned study, the duration of therapy, and the fact that three out of seven initially tested dogs were revealed to be receiving a placebo once the trial was unblinded. Blood samples were transferred to blood spot cards and then stored at −80°C for multiple months prior to analysis, which could possibly have impacted the reported rapamycin concentrations. However, blood spot cards are a highly stable form of storage, with some studies utilizing cards stored for over a decade.[54] In regard to the analysis, measurements below 1 ng/ml were obtained and reported, despite the fact that these measurements fall below the formal limit of detection for the assay, which is a statistical bound on confidence that the measured value is different from zero. We feel comfortable including these low measurements because all of the dogs receiving rapamycin had positive measurement values, while none of the dogs receiving placebo had non-zero measurement values for any of the timepoints. We recognize, however, that the low measured values may contribute increased variance and higher error.

Finally, even though all dogs were offered food with their treatment administration, only one dog (#4) ate. Coincidentally, or perhaps consequently, this participant also had the highest blood levels of rapamycin and longest blood half-life. Rapamycin pharmacokinetics in humans have previously been shown to be impacted by the consumption of food, with a high-fat meal resulting in reductions in C_max_ but increases in AUC by 23 to 35%.[49] This, however, has not been investigated in dogs.[49] Given this, future investigation into whether food consumption does affect rapamycin absorption in dogs would be valuable, as this might influence the protocol of the upcoming long-term low-dose study, and other uses of rapamycin in this species, by recommending treatment administration at a specified time relative to a meal.

## Conclusions

The results of this study indicate that low-dose (0.025 mg/kg) oral rapamycin, administered three times a week for one to five months to healthy, middle-aged large breed dogs, resulted in detectable concentrations of rapamycin in four dogs that was measurable in ng/ml. While the long-term benefits and ideal dosing schedule of rapamycin are still not known, this study demonstrates that low doses produce notable systemic exposure and that intermittent treatment may allow clearance of rapamycin from blood prior to the next dose.

## Abbreviations

HPLC/MS/MS: High performance liquid chromatography – mass spectroscopy

## Author Contributions

Conceptualization – Daniel E.L. Promislow, Matt Kaeberlein, and Kate E. Creevy

Funding Acquisition – Daniel E.L. Promislow, Matt Kaeberlein, and Kate E. Creevy

Investigation – Jeremy B. Evans and Ashley J Morrison

Methodology – Daniel E.L. Promislow, Matt Kaeberlein, and Kate E. Creevy

Project Administration – Kate E. Creevy

Resources – Kate E. Creevy, Marisa Lopez-Cruzan, and Martin A. Javors

Validation – Marisa Lopez-Cruzan and Martin A. Javors

Writing – Original Draft Preparation – Jeremy B. Evans and Ashley J Morrison

Writing – Review & Editing – Daniel E.L. Promislow, Matt Kaeberlein, Kate E. Creevy, Marisa Lopez-Cruzan, and Martin A. Javors

## Acknowledgments

This research was supported by the William H. Donner Foundation and by grant U19AG057377 from the National Institutes on Aging. We are also grateful for the assistance of Drs. Virginia Fajt and Madeline Droog with the PK analysis. The content is solely the responsibility of the authors and does not necessarily represent the official views of either funding body. Funding sources did not have any involvement in the study design, data analysis and interpretation, or writing and publication of the manuscript.

